# Evolution of dispersal can rescue populations from expansion load

**DOI:** 10.1101/483883

**Authors:** Stephan Peischl, Kimberly J. Gilbert

## Abstract

Understanding the causes and consequences of range expansions or range shifts has a long history in evolutionary biology. Recent theoretical, experimental, and empirical work has identified two particularly interesting phenomena in the context of species range expansions: (i) gene surfing and the relaxation of natural selection, and (ii) spatial sorting. The former can lead to an accumulation of deleterious mutations at range edges, causing an expansion load and slowing down expansion. The latter can create gradients in dispersal-related traits along the expansion axis and cause an acceleration of expansion. We present a theoretical framework that treats spatial sorting and gene surfing as spatial versions of natural selection and genetic drift, respectively. This model allows us to study analytically how gene surfing and spatial sorting interact, and to derive the probability of fixation of pleiotropic mutations at the expansion front. We use our results to predict the co-evolution of mean fitness and dispersal rates, taking into account the effects of random genetic drift, natural selection and spatial sorting, as well as correlations between fitness- and dispersal-related traits. We identify a “rescue effect” of spatial sorting, where the evolution of higher dispersal rates at the leading edge rescues the population from incurring expansion load.

## Introduction

Understanding the demographic, ecological, and evolutionary forces that determine the evolution of a species range has been a central area of research since the early days of evolutionary biology (Darwin, 1859; Sexton et al., 2009). Over the last decade, the fact that species range expansions impact multiple evolutionary and ecological processes in peripheral populations has been thrown into the spotlight both theoretically and empirically (see e.g., Bosshard et al., 2017; Brown et al., 2013; Burton et al., 2010; Fronhofer and Altermatt, 2015; González-Martínez et al., 2017; Hallatschek and Nelson, 2010; Klopfstein et al., 2006; Peischl et al., 2013; Shine et al., 2011; Travis et al., 2007; Van Dyken et al., 2013; Weiss-Lehman et al., 2017). This shift in thinking about the dynamic processes forming species ranges has led to the observation that evolutionary and ecological dynamics at the front of a range expansion can differ considerably from those in the core of a species range. The set of traits that allow a species to colonize and expand its range might thus be very different from those that allow a species to successfully persist in new habitat. In this work, we study the co-evolution of two traits that are highly relevant in the context of species range expansion, namely an individual’s fitness and its dispersal abilities.

A first key process in determining evolutionary processes during a range expansion is genetic drift. In a seminal paper, Edmonds et al. (2004) showed that strong genetic drift at the front of range expansions can lead to the rapid increase of random neutral variants along the expansion axis, a process now known as gene surfing (Klopfstein et al., 2006). Gene surfing also affects selected variants (Travis et al., 2007) and can lead to an accumulation of deleterious mutations in marginal populations (Hallatschek and Nelson, 2010). This accumulation of deleterious mutations has been been termed expansion load and has been the subject of several theoretical (Gilbert et al., 2017; Peischl et al., 2013; Peischl and Excoffier, 2015; Peischl et al., 2015), experimental (Bosshard et al., 2017; Weiss-Lehman et al., 2017), and empirical studies (González-Martínez et al., 2017; Henn et al., 2016; Peischl et al., 2018; Willi et al., 2018). Expansion load stems from the repeated founder events at expanding wave fronts that reduce the efficiency of selection which would otherwise purge most incoming deleterious mutations. In this sense, the evolutionary dynamics at the front of expanding populations are similar to that of mutation accumulation experiments (Bosshard et al., 2017). Several factors contribute to the dynamics and severity of expansion load. Theoretical work has identified that fast-growing species with low dispersal rates are most likely to accumulate harmful mutations (Peischl et al., 2013). The distribution of fitness effects and the degree of dominance of mutations also have a strong impact on the evolution of expansion load (Gilbert et al., 2018; Peischl et al., 2013; Peischl and Excoffier, 2015).

A second important process that can arise during range expansions is the evolution of dispersal-related traits (see, e.g., Bouin and Calvez, 2014; Deforet et al., 2017; Phillips and Perkins, 2017; Phillips et al., 2006; Simmons and Thomas, 2004; Travis and Dytham, 2002), which has been termed spatial sorting (Shine et al., 2011). When a population possesses heritable variation in dispersal abilities, colonists at the range front result disproportionately from individuals with greater dispersal propensity. Individuals are thus sorted over space according to their dispersal abilities, with more dispersive individuals at the range edge, similar to the increase in frequency of beneficial mutations over time due to natural selection (Phillips and Perkins, 2017; Shine et al., 2011). Spatial sorting thus increases dispersal propensity at the front as these individuals mate assortatively, potentially accelerating the speed of a range expansion (Burton et al., 2010; Cwynar and MacDonald, 1987; Hughes et al., 2003; Phillips et al., 2008; Travis et al., 2007). Spatial sorting has most notably been described in the invasive expansion of cane toads (*Rhinella marina*) across Australia (Phillips et al., 2006), but has been observed in several other systems (Fronhofer and Altermatt, 2015; Simmons and Thomas, 2004; Van Ditmarsch et al., 2013; Weiss-Lehman et al., 2017).

A few theoretical studies have focused on the co-evolution of fitness- and dispersal-related traits during range expansions. Using individual-based simulations, Burton et al. (2010) studied the evolution of resource allocation for three life-history traits during range expansions: dispersal, reproduction, and competitive ability. They found that dispersal and reproductive abilities generally increase on the expansion front, whereas competitive abilities decrease as compared to the core. Using a deterministic serial founder effect model with discrete demes, Phillips and Perkins (2017) showed that a mutation that alters both fitness and dispersal abilities will be positively selected on an expansion front if the product of migration rate and fitness is greater than that of an individual with the wild-type allele. Deforet et al. (2017) study the evolution of expansion speed using a deterministic reaction-diffusion type model in continuous space, finding that a mutation can invade the expansion front if it leads to an increase in expansion speed. The expansion speed in their model is proportional to the square root of the product of migration rate and growth rate, and hence any mutation that increases the product of migration rate and growth rate will be positively selected at the expansion front. Despite modelling differences, the conclusions of Phillips and Perkins (2017) and Deforet et al. (2017) are strikingly similar, in the sense that the product of fitness (or a fitness-related trait such as growth rate) and dispersal rates is what determines whether a mutation is adaptive for expansion or not. The reason for their similar conclusions is that both studies focus on a deterministic model with two key aspects: the ability of reaching the front (determined by dispersal rates) and the chance of surviving on the front (determined by fitness or growth rates). It remains unclear, however, how genetic drift, mutation rates, correlations between traits, and the relationship between fitness, growth rates and expansion speed may influence evolutionary dynamics at expansion fronts.

There is striking evidence for both spatial sorting and expansion load from experimental evolution studies. Using *Escherichia coli*, Bosshard et al. (2017) has shown that fitness decreased during expansion on agar plates due to a random accumulation of new incoming mutations. Intriguingly, there are signals for an increase in expansion speed during early phases of the experiment, potentially due to loss of function in genes related to flagella production, which might allow bacteria to reach the expansion front more easily (Bosshard et al., 2018). However, in the long term, expansion speed was found to decrease over time due to reduced growth rates and competitive abilities, corroborating theoretical results (Peischl et al., 2015). Van Ditmarsch et al. (2013) performed similar experiments with *Pseudomonas aeruginosa* where they found strong signals of convergent evolution of a “hyperswarming” phenotype with increased numbers of flagella per individual. Even though growth rates in the evolved strains were lower as compared to the wild-type, the expanded populations out-competed ancestral populations, seemingly due to their increased dispersal abilities (Deforet et al., 2014). In addition to using different species, another key difference between these two experimental studies is the viscosity of the agar environment, and hence the mechanisms of dispersal in the bacteria. While Bosshard et al. (2017) used solid agar (at a concentration of 1.5% (w/v)) where bacteria are “pushed” to the front, Van Ditmarsch et al. (2013) used soft agar (at a concentration of 0.3% (w/v)) that allowed for active dispersal of bacteria via swarming. The extent to which these differences have contributed to the different outcomes of the two experiments remains unclear. These examples of disparate outcomes for evolution of dispersal and fitness emphasize the need to fully understand the theoretical underpinnings of expansion load and spatial sorting and to identify when they may complement or disrupt each other.

In this study, we derive theoretical expectations for when and how interactions between genetic drift, natural selection, and spatial sorting may unfold. Our framework allows a detailed analytic treatment and can be used to predict the co-evolutionary dynamics at expansion fronts. A key analytic result is the derivation of the fixation probability of a pleiotropic mutation affecting both fitness and dispersal-related traits.

## Model and Results

We model the evolutionary dynamics of allele frequencies at the front of a one-dimensional range expansion, combining the approaches of Peischl et al. (2013, 2015); Phillips and Perkins (2017); Slatkin and Excoffier (2012). Consider an infinite stepping-stone model of demes, labelled *d* = 1, 2, 3, …, *n*. The carrying capacity of each deme is denoted *K*. Initially, only a subset of demes is colonized, and all other demes are empty. *d* _*f*_ (*t*) will denote the most recently colonized deme at time *t*, which we call the expansion front. Individuals are haploid, and we consider a single locus with two alleles denoted *a* and *A.*These alleles can affect either fitness or dispersal rates, or both. Let *p* denote the frequency of the mutant allele *A* at the expansion front, that is, in deme *d* _*f*_. Note that the dependence on *t* is omitted for the sake of simplicity. The fitness of wild-type and mutant alleles are denoted *w*_*a*_ and *w*_*A*_, respectively, and the selection coefficient *s* of the mutant allele *A* is given by 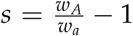. During the dispersal phase, wild-type individuals migrate to neighboring demes with probability *m*_*a*_ and mutants with probability *m*_*A*_. Analogous to the selection coefficient *s*, we define the effect on dispersal rate from a mutant allele as 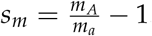.

A key simplifying assumption in our model is that we model the colonization of new demes as discrete founder events occurring every *T* generations (see e.g., Peischl et al., 2013, 2015). When a deme is at carrying capacity, a propagule of size *F* is placed into the next empty deme *d* _*f*_ (*t*)+ 1. The population then grows exponentially for *T* generations until the new deme’s carrying capacity is reached. The size of the propagule is determined by the dispersal abilities of individuals at the expansion front. Let 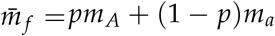 denote the average migration rate in the population. The size of the propagule is then 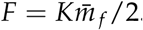. The factor 1/2 is due to the fact that individuals migrate to each of the two neighboring demes with the same probability. During the growth phase, migration is ignored. Assuming exponential growth at rate *r* = log(*R*), this yields 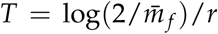 (Peischl et al., 2013). This model is a good approximation to range expansions with continuous gene flow when growth rates are larger than migration rates (Peischl et al., 2013). We also consider the limiting case where *r* is so large that a deme grows to carrying capacity within a single generation *T* = 1, independently of the number of founders *F*. Figure 1 shows a sketch of the model that illustrates how mutations can be positively selected on expanding wave fronts based on either an increase in migration rates (Figure 1A) or an increase in relative fitness (Figure 1B).

**Figure 1:**
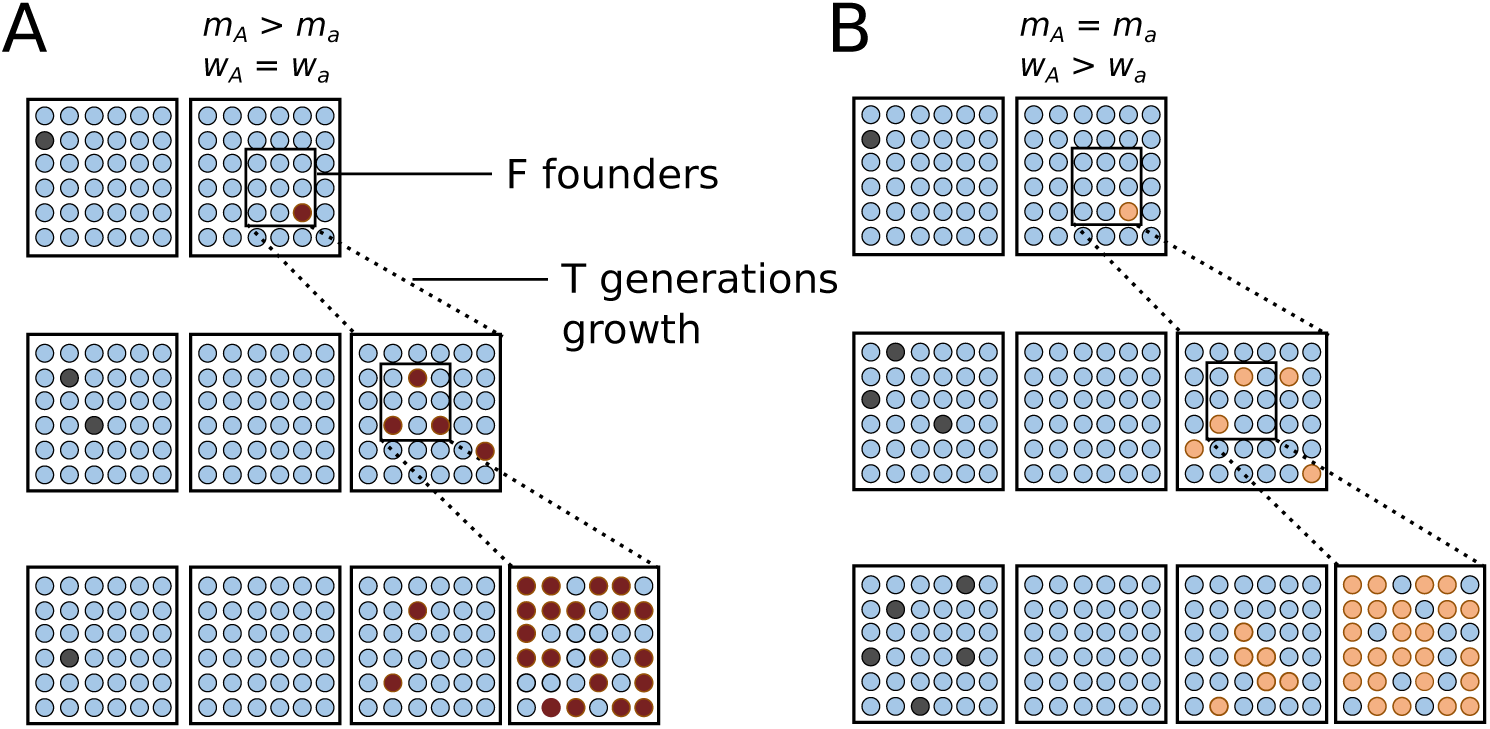
Sketch of the model. A: a mutation with higher migration rate (red) but same fitness as wild-type individuals (blue). The mutation can increase in frequency at the expansion front because it is more likely to be among the *F* founders as compared to wild-type mutations. B: a mutation with higher fitness than the wild type, but with same migration rate. The mutation (orange) has the same probability to be among the *F* founders as the wild type (blue), but it can spread at the expansion front due to higher reproductive success during the *T* generations of growth during which natural selection acts. In both panels, dark gray circles show the evolution of an equivalent mutation in the core of the species range for comparison.

### Fixation of new mutations

We show in Appendix A that the probability of fixation of a mutation with initial frequency *p*_0_ at the expansion front is given by

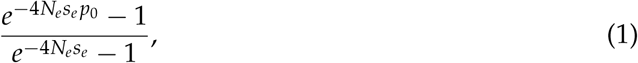

where we define an effective selection coefficient *s*_*e*_ = *sT* + *s*_*m*_ and an effective population size *N*_*e*_ = *F*. Equation (1) shows that mutations can be under positive selection at the expansion front for two reasons: (i) increasing an individual’s fitness (*s >* 0) or (ii) increasing the migration rate (*s*_*m*_ *>* 0). If *s*_*m*_ = 0, we recover the fixation probability on the expansion front derived in Peischl et al. (2013). Then, natural selection is most efficient when *R* is small (Figure 2B) and 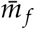 is large (Peischl et al., 2013, Figure S4B). If *s* = 0, the fixation probability of a mutation modifying the dispersal probability by a factor of 1 + *s*_*m*_ is equivalent to that of a mutation with selective advantage *s*_*m*_ in a stationary population of size *F* (Kimura, 1962). This shows that spatial sorting can indeed be viewed as an analog to natural selection across space as proposed by simulation studies (Shine et al., 2011) and deterministic models (Phillips and Perkins, 2017). Note that our model can be seen as a stochastic version of the model presented in Phillips and Perkins (2017) if *T* = 1, i.e., if a new deme is colonized each generation.

**Figure 2:**
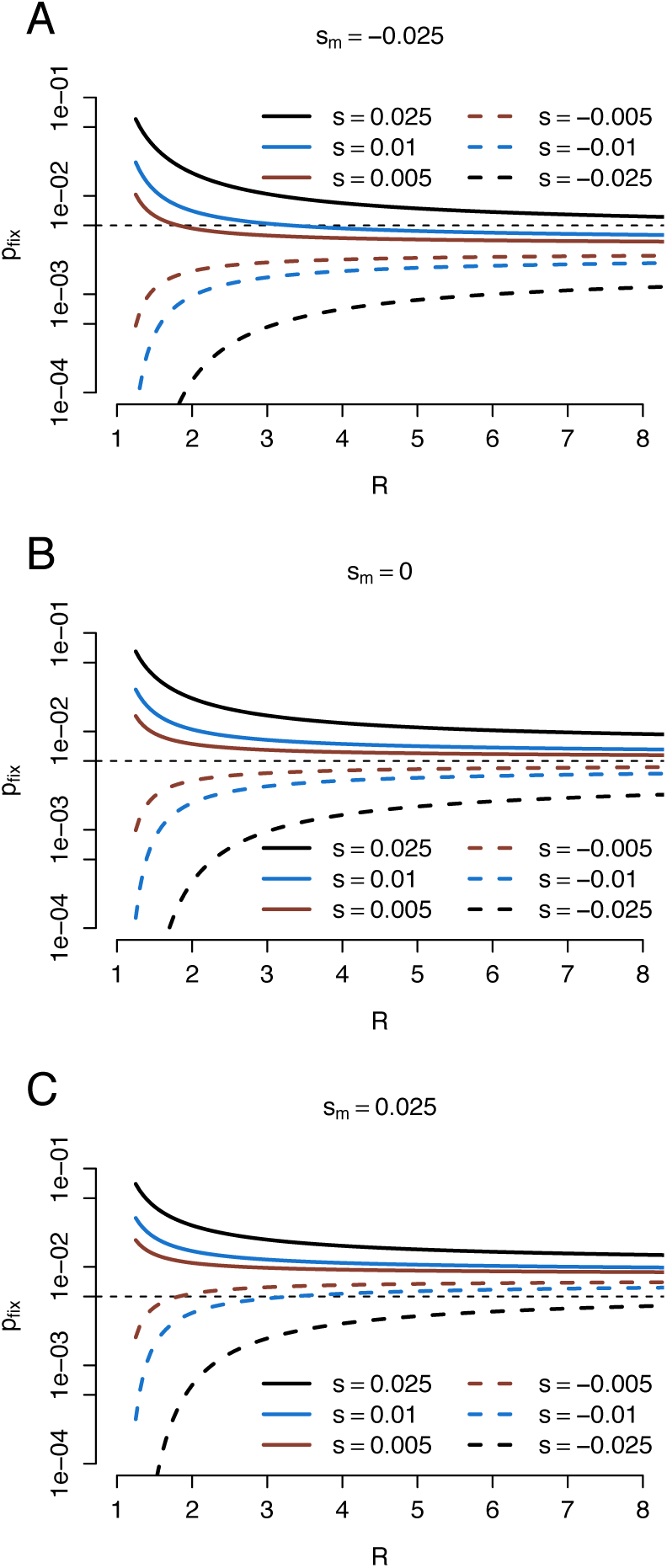
Fixation probability of pleiotropic mutations as a function of population growth rate. Dashed lines indicate deleterious mutations (negative selection coefficient, *s*) while solid lines indicate beneficial mutations.

In the following we denote mutations with *s*_*e*_ *>* 0 as adaptive for expansion, since they can spread at the front because of the joint actions of natural selection and spatial sorting. We refer to mutations with *s*_*e*_ *<* 0 as maladaptive for expansion since they can only establish via genetic drift. Equation (1) shows that natural selection is most efficient if 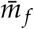 is large and *R* is small (see also Peischl et al. 2013). Likewise, spatial sorting is most efficient if 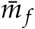 is large because drift during founder events decreases with increasing migration rates (Figure S2). The growth rate *R* has no impact on the fixation probability if *s* = 0 (Figure S1), since it only affects the length of the growth phase during which natural selection acts but not the number of founders, *F*, or the probabilities of individuals to migrate to a new deme.

### Pleiotropic mutations

We next consider mutations that affect both the fitness as well as the dispersal ability of a carrier. As expected, mutations that increase both fitness and migration rates (*s, s*_*m*_ *>* 0) are always positively selected (solid lines in Figure 2C) and mutations with *s, s*_*m*_ *<* 0 are always negatively selected at the expansion front (dashed lines in Figure 2A). In both cases, the efficacy of selection for expansion decreases with increasing growth rate *R* (Figure S2) because the time *T* during which natural selection can act becomes shorter.

If there is a trade-off between fitness- and dispersal-related traits such that *s*_*m*_ *<* 0 *< s* or *s <* 0 *< s*_*m*_, the growth rate of the population, *R*, affects the strength as well as the direction of selection for a given mutation (Figure 2). In general, if growth rates are low, natural selection is more effective than spatial sorting because of the longer periods, *T*, between consecutive founder events during which selection can act (Figure 2), whereas spatial sorting is only acting during the sampling of new founders (Figure 1). Thus, for low *R*, fixation probabilities are close to that of mutations with effect *s* in stationary populations of size *F*. On the other hand, if *R* is large such that *T* is close to 1, both spatial sorting and natural selection contribute equally to the fixation probability (Figure 2), which is then similar to a mutation with effect *s* + *s*_*m*_ in a stationary population of size *F*. We find that a mutation with *s >* 0 *> s*_*m*_ has a higher fixation probability as compared to a neutral mutation (*s* = *s*_*m*_ = 0) if *R < m*^*s*/*s*_*m*_^ (Figure 2A), and a mutation with *s <* 0 *< s*_*m*_ has a higher fixation probability if *R > m*^*s*/*s*_*m*_^. Taken together, this means that spatial sorting and expansion load should more readily impact populations with high growth rates, especially if increasing dispersal rates is costly in terms of fitness.

Figure 3 illustrates the effect of the average migration rate at the expansion front on the fixation probability of mutations. For very small values of 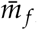, the number of founders is close to *F* = 1 and selection for expansion is therefore virtually absent (as in mutation accumulation experiments). The fixation probability is then close to that of a neutral mutation (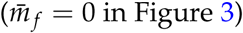 in Figure 3). As 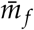 increases (to ≈ 0.1 in Figure 3) the fixation probability of a pleiotropic mutation is driven more by the action of natural selection rather than the action of spatial sorting. This is when *T* is sufficiently large to allow the contribution of natural selection to outweigh that of spatial sorting in the effective selection coefficient *s*_*e*_. For even larger values of 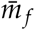, the time between founder events will decrease as propagule size increases and eventually approach *T* = 1 such that *s* and *s*_*m*_ will contribute equally to the fixation probability. Thus, if *s* + *s*_*m*_ *>* 0 and *s <* 0 *< s*_*m*_ (Figure 3A) or if *s* + *s*_*m*_ *<* 0 and *s*_*m*_ *<* 0 *< s* (Figure 3B), the direction of selection for pleiotropic mutations may change with increasing 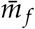.A mutation with *s <* 0 *< s*_*m*_ has a higher fixation probability as compared to a neutral mutation (*s* = *s*_*m*_ = 0) if 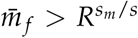. Conversely, pleitropic mutations with *s >* 0 and *s*_*m*_ *<* 0 will have a higher fixation probability as compared to neutral mutations if 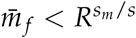.

**Figure 3:**
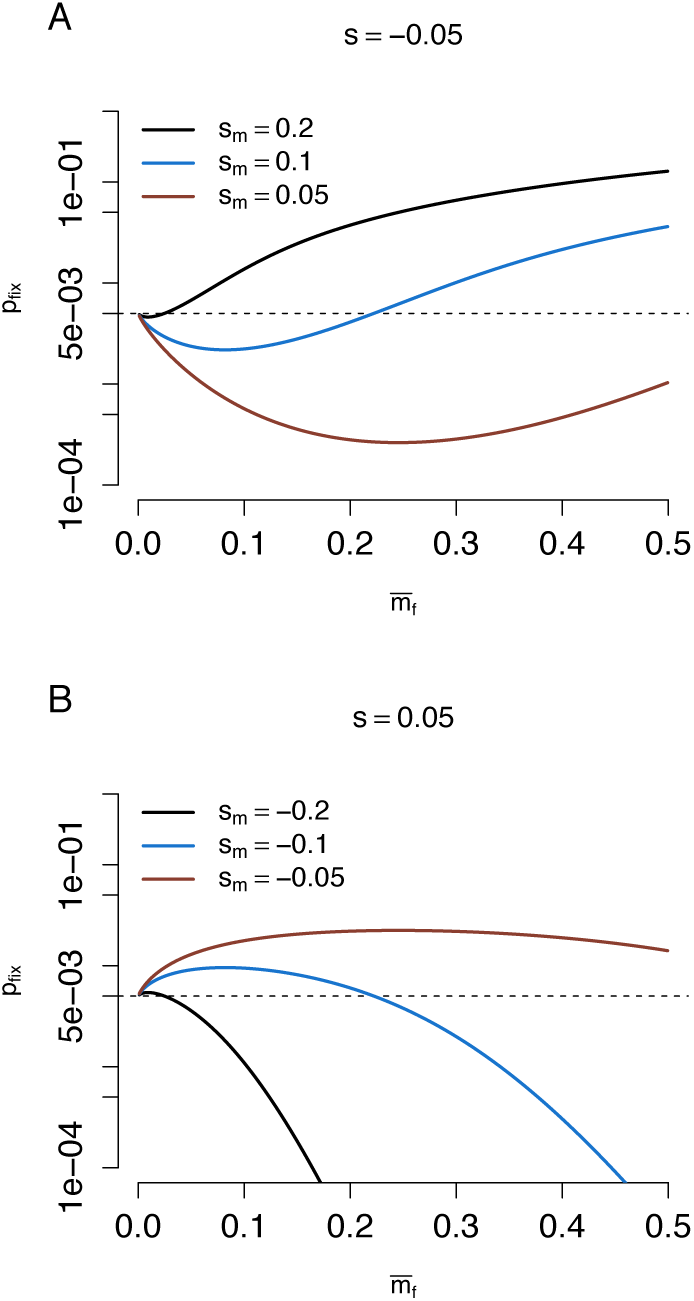
Fixation probability of pleiotropic mutations as a function of mean migration rate at the front. *s* is the selection coefficient for fitness-impacting mutations while *s*_*m*_ is the selection coefficient for dispersal-impacting mutations.

### Co-evolutionary dynamics

We next study the co-evolution of mean fitness and migration rates in expanding populations taking into account the interactions of mutation rates, the distribution of fitness effects (DFE) of new mutations, and genetic correlations in fitness and migration-related traits. In the following we assume that selection is soft such that population mean fitness does not affect growth rates or carrying capacities. Consequently the parameters *T* and *F* are independent of the evolution of mean fitness (cf. Peischl et al., 2015), and following equation (1), the evolution of mean fitness does not impact the evolution of migration modifiers. However, the amount of migration into new empty demes affects both the parameters *F* and *T* (Peischl et al., 2015), which in turn determine the efficacy of selection and the strength of drift at the expansion front. We approximate the evolution of mean fitness and migration rate, analogous to the model in Peischl et al. (2015), and consider mutations that can affect both migration rates and fitness simultaneously. Let *u*(*s, s*_*m*_) denote the mutation rate of mutations with effect *s* on fitness and *s*_*m*_ on the migration rate. We assume that *s* and *s*_*m*_ are drawn from a bi-variate distribution with mean 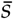 and 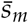, variance *V*_*s*_ and *V*_*m*_, and correlation *ρ*. Appendix B shows that we can approximate the dynamics of mean fitness and migration rate at the front by

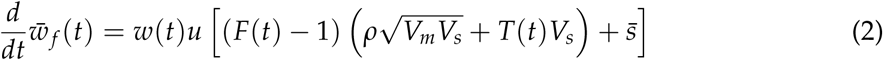

and

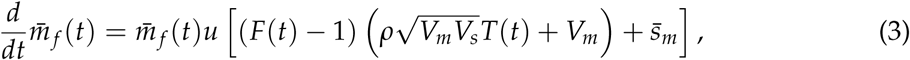

where 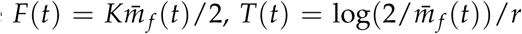 and *u* is rate at which new mutations occur per individual and generation.

In general, the mean mutational effect of mutations affecting fitness will be negative 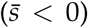 as most new incoming mutations are deleterious (Eyre-Walker and Keightley, 2007). Thus, expansion load will generally occur unless one of the following is true: the variance of the distribution of fitness *V*_*s*_ is sufficiently large (thus increasing the proportion of beneficial variants in the DFE, see also Peischl et al. 2013), the covariance of a mutation’s fitness effects with effects on migration related traits is positive and sufficiently large, or the carrying capacity of demes at the expansion front is sufficiently large. A negative correlation between fitness-related and migration-related traits can increase the chance for expansion load to occur.

Spatial sorting is expected to occur if there are sufficiently many new mutations that increase migration rates (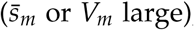 or *V*_*m*_ large), if mutations that increase migration rates also increase fitness (*ρ >* 0), or if population size is sufficiently large. If fitnessand migration-related traits are independent, that is *ρ* = 0, then 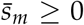 implies that migration rates will always increase at the expansion front. If there is, however, a trade-off between migrationand fitness-related traits such that *ρ <* 0, the average migration rate at the front can decrease despite 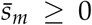. Thus, if an increase in migration is costly in terms of fitness, expansion load can constrain spatial sorting. A positive correlation between effects on migration and fitness (*ρ >* 0) will generally increase the chance for spatial sorting to occur, as well as reduce the chance for expansion load to accumulate.

### Evolution of dispersal can rescue expanding populations

While a detailed analytic analysis of eqs. (2) and (3) is mathematically challenging and beyond the scope of this paper, we can gain some intuition from the case when growth rate is strong such that newly colonized demes reach carrying capacity within a single generation (*T* = 1). This is usually the case when *r >>* 1. Here, we find that 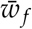 increases over time if

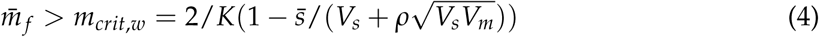

and *m*_*f*_ increases over time if

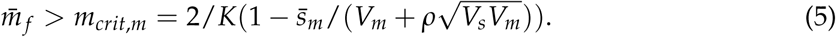

Figure 4 illustrates the dynamics of mean fitness and migration rate at the expansion front as given by eqs. (2) and (3) (arrows in Figure 4) and compares these dynamics with the outcome of individual-based simulations in a serial founder effect model (as depicted in Figure 1). Even though eqs. (2) and (3) can be solved analytically, we proceed by describing the dynamics using a geometric approach that allows us to exhaustively identify all qualitatively different evolutionary regimes. If 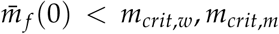, an expanding population will not evolve increased dispersal and will also suffer from expansion load (green lines in Figure 4). On the other hand, if 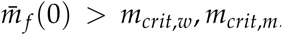, both mean fitness and the average migration rate at the expansion front will increase (red lines in Figure 4). More interestingly, if 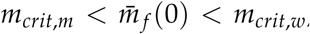, expansion load will accumulate and migration rates will also increase over time. Thus, we eventually observe a “rescue effect” when 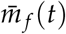 surpasses *m*_*crit,w*_, in the sense that founder events become less drastic and selection at the expansion front becomes sufficiently efficient so that mean fitness will start to increase over time (see blue lines in Figure 4).

**Figure 4:**
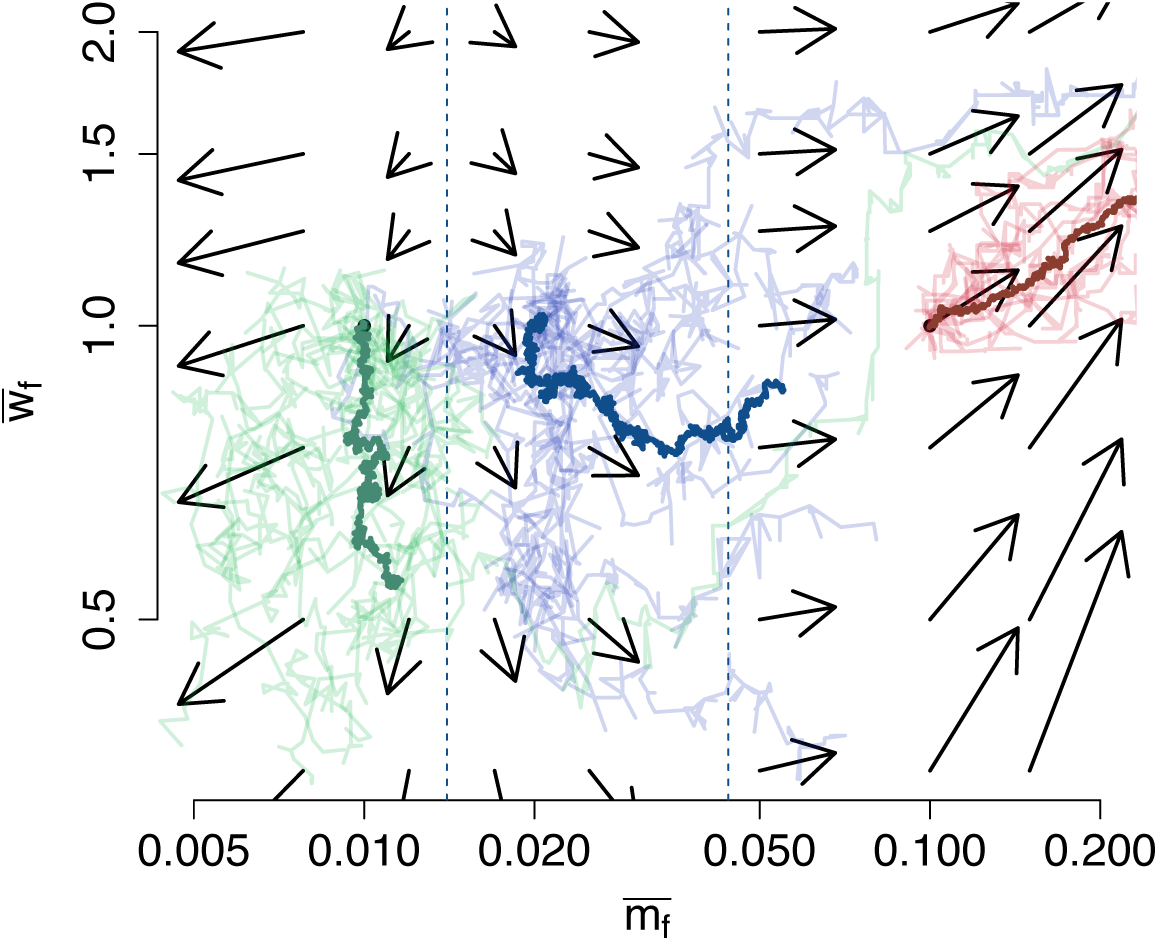
Individual-based simulations of the co-evolution of mean fitness and migration rate in a population undergoing serial founder events, as depicted in Figure 1. The arrows show the vector field generated by the differential equations (2) and (3), and indicate the direction of evolution as predicted by the analytic theory. The thin lines show the outcome of single simulation runs. The thick lines show the average across 10 simulation runs for each initial condition. The different colors correspond to different initial migration rates. Mutations occur at rate *u* = 0.01 per individual per generation and their effects are drawn from a bi-variate Gaussian distribution with parameters 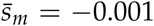, 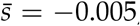, *V*_*m*_ = 0.004, *V*_*s*_ = 0.002 and *ρ* = 0. The remaining parameters are *K* = 500, 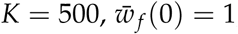, and 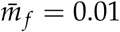 (green), 0.02 (blue) and 0.1 (red).

### Individual-based range expansion simulations

Using an individual-based model first developed in Peischl et al. (2013), we simulate populations undergoing range expansions with both the evolution of dispersal and fitness. The key difference to the serial founder effect model in Figure 4 is that we simulate the whole species range instead of just the deme at the leading edge, and that gene flow occurs every generation rather than just during colonization events. In particular, we model a linear, 1-dimensional discrete landscape of 1000 demes with a stepping-stone migration model. Each deme has a carrying capacity of *K* = 1000 and an initial migration rate *m* = 0.01. The 5 left-most demes are initiated at carrying capacity and burned in for 6000 generations, after which free expansion into subsequent empty demes is allowed. Individuals each have the potential to accumulate deleterious load through 1300 bi-allelic, unconditionally deleterious loci, or increase fitness from 700 bi-allelic, unconditionally beneficial loci. Fitness is multiplicative across these 2000 loci with genomewide mutation rate *U* = 0.2 and an equivalent potential for back-mutations to the wild type. Fitness effects are fixed at *±*0.01 and are additive (heterozygote fitness is perfectly intermediate to homozygotes). Generations are non-overlapping and growth is instantaneous in newly-colonized demes. The dispersal trait is modelled as a quantitative trait such that each individual inherits its migration rate from as the average of both parents’ trait value plus a random mutational deviation drawn from a Normal distribution with mean 0 and standard deviation 0.005 or 0.01 for either a low or high rate of dispersal evolution, respectively. Migration rate is constrained between 0 *< m <* 0.5.

For these parameter values, equations (4) and (5) indicate that we expect an increase in dispersal rates independently of initial conditions, and that expansion load occurs if 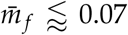 and ceases to occur if 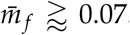. In agreement with these predictions, we find that mutation load does accumulate during expansion as a result of gene surfing of deleterious mutations, but also that as dispersal evolves, spatial sorting leads to the rescue of fitness at the range front (Figure 5). The rescue effect is particularly strong under a higher rate of dispersal evolution (Figure 5C), where migration rate evolves to be close to 0.5. Under both low and high rates of dispersal evolution, fitness loss is reduced and then reversed, an effect opposite to that expected for fast range expansions in the absence of dispersal evolution (Gilbert et al., 2017; Hallatschek and Nelson, 2010; Peischl et al., 2013, 2015). Conversely, in the absence of the evolution of dispersal, fitness is continually lost throughout the course of expansion.

**Figure 5:**
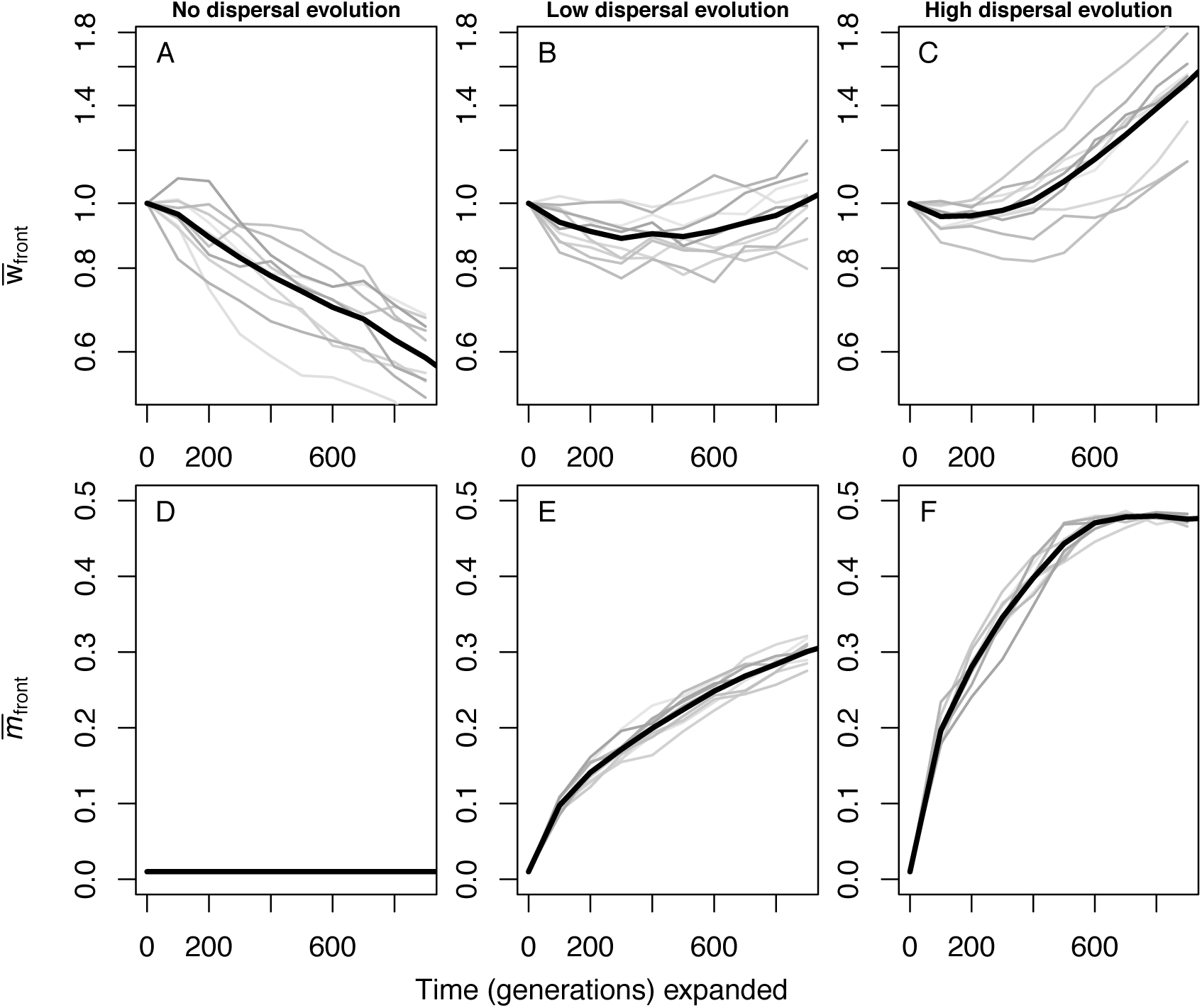
Fitness (A-C) or migration rate (D-F) measured at the front edge of a range expansion either in the absence of the evolution of migration or a low (standard deviation of mutational effect 0.005) or high (standard deviation of mutational effect 0.01) rate of evolution of migration. Individual replicate simulations are shown in gray while the mean is shown by the thick black line. Starting fitness is scaled to 1 for comparison, and all other parameters are as described in text.

## Discussion

The question of what makes an organism successful at colonizing new habitats is highly relevant in evolutionary biology, (Sexton et al., 2009), conservation biology (e.g., for predicting invasiveness of species, Pejchar and Mooney 2009), and evolutionary medicine (e.g., in the context of cancer growth, Waclaw et al. 2015). In this work, we present an analytically tractable theoretical framework for the co-evolution of fitnessand dispersalrelated traits that builds upon classical models in population genetics. We show that a given mutation with pleiotropic effects can be positively or negatively selected at the expansion front, depending on the current growth rate and migration rate at the expansion front (see Figures 2 and 3). Furthermore, we show that while migration rates and growth rates both affect the expansion speed in similar ways, their effect on the strength and direction of selection at the expansion front can be quite different in finite populations (see Figures 2 and 3). Finally, we used our results to predict the co-evolutionary dynamics of fitness and dispersal during range expansions (see equations (2) and (3), and Figure 4). For the special case of high growth rates and soft selection, our model allowed us to exhaustively characterize the co-evolutionary dynamics, and to identify conditions when the evolution of dispersal can rescue a population from expansion load (see equations (4) and (5), and Figure 4). Individual-based simulations of range expansions confirmed our analytic results (see Figure 5).

Our study generalizes the results of Phillips and Perkins (2017), who studied the co-evolution of dispersal and fitness during range expansions with a constant speed of 1 deme per generation (*T* = 1) using a deterministic model similar to ours. As expected, in the case of infinite population size and *T* = 1, our results are in perfect agreement with those of Phillips and Perkins (2017), meaning that mutations are adaptive for expansion if and only if *s*_*e*_ = *s* + *s*_*m*_ *>* 0, which, for small *s* and *s*_*m*_ is equivalent to the conditions derived in Phillips and Perkins (2017). We note that this condition for invasion of new mutations is also equivalent to the condition found by Deforet et al. (2017) in a deterministic continuous space model if one treats growth rate in their model as equivalent to fitness. It would be interesting to see whether and how results from deterministic continuous space models further generalize to finite populations, and to better understand the role of genetic correlations on spatial sorting and expansion load. A direct comparison of our results with those of Deforet et al. (2017) is difficult because their assumptions differ regarding the interplay of fitness, growth rates, and gene flow in the modelling approaches.

The simplicity of our model comes at a cost as we made several simplifying assumptions. Perhaps most importantly, we employ a separation of time scales argument that allows us to model evolution of the leading edge population independently from the core. We have previously shown that this is a good approximation to models with continuous gene flow between demes as long as the growth rate of the population is sufficiently large (Peischl et al., 2013). We thus expect our results to be valid even if dispersal rates are large, as long as growth rates are even larger (see Figure 5 for results from individual-based simulations). If growth rates are on the order of dispersal rates or lower, we expect our model to underestimate the strength of drift because gene flow will lead to a more gradual decrease in population size towards the expansion front (c.f. Hallatschek and Nelson, 2010). Therefore, the rescue effect we identified with our model might be less relevant for species with growth rates and migration rates of similar magnitudes. In this case, a more suitable modelling approach would be assuming continuous space, e.g., using reaction-diffusion equations as in Deforet et al. (2017). Including the effects of genetic drift, however, is somewhat harder in continuous-space models (but see e.g. Barton et al. 2013; Brunet and Derrida 2001; Hallatschek 2011).

We focused on expansion along a one-dimensional habitat. This should be a good approximation for range expansion along a narrow two-dimensional corridor (Peischl et al., 2013). In wider habitats, the evolutionary dynamics at the expansion front might be quite different from what is expected in the one-dimensional case (see e.g., Polechová and Barton, 2015). In particular, lateral gene flow perpendicular to the axis of expansion can restore genetic diversity and hence prevent some of the negative consequences of increased drift at the expansion front (Peischl et al., 2013). Previous studies have shown that a wider expansion front can reduce the rate at which expansion load is built up and lead to faster recovery after the expansion (Gilbert et al., 2018). With spatial sorting it remains unclear how a two-dimensional expansion front would affect the outcome. Gene flow might have very different effects on expansion speed and genetic diversity, depending on its direction relative to the expansion axis. One might thus expect that not only the rate or distance of dispersal evolves, but also the direction of dispersal (see e.g., Lindström et al., 2013).

For the sake of simplicity we assumed haploid individuals, but our model can be readily extended to sexually reproducing, diploid organisms (see e.g., Phillips and Perkins 2017). Since the evolutionary dynamics of diploid and haploid individuals are equivalent in the case of co-dominant (multiplicative) mutations (Bürger, 2000), our model can be applied directly to diploid individuals. Additionally, while it would be straightforward to include dominance in our model (Gilbert et al., 2018; Peischl and Excoffier, 2015), adding epistatic interactions across loci would be much more difficult. Furthermore, we ignored the effects of clonal interference in the derivation of equations (1), which could lead to an overestimation of the fixation probability of beneficial mutations. Our results should be good approximations if recombination is strong or if mutations occur infrequently so that multiple segregating mutations rarely interact (that is, if *uK <* 1). However, because mutations are either fixed or lost very quickly at expanding fronts, we expect our results to hold even if mutation rates are fairly high such that *uK >* 1 (see Figure 4).

We assumed that selection is soft, i.e. local carrying capacities and growth rates do not depend on population mean fitness. The co-evolutionary dynamics under hard selection might be somewhat different from those in our our model (see e.g., Peischl et al., 2015) since growth rates and carrying capacities affect expansion speed (Skellam, 1951), the amount of genetic drift, and the efficacy of spatial sorting and natural selection at the expansion front (Hallatschek and Nelson, 2010). In particular, while increasing growth rates render natural selection at expansion fronts less efficient due to the reduced time between subsequent colonization events (see Figure 2), the efficacy of natural selection increases with increasing dispersal (see Figure 3). A model with hard selection would thus lead to additional feedback between evolutionary processes at the expansion front and the efficacy of natural selection and spatial sorting.

We have presented a theoretical study that shows how the evolution of dispersal can serve as a factor to reduce or even eliminate expansion load. To further test our model in experimental or empirical settings, comparing fitness evolution during geographic spread in tandem with dispersal evolution will prove especially illuminating. This is most approachable in experimental evolution studies where understanding these trajectories simultaneously will inform how often dispersal is positively or negatively correlated with changes in fitness. This may also provide insights into understanding the distribution of fitness effects for new mutations and for dispersal-impacting mutations by fitting a model to data from an experimental study (e.g., as in Bosshard et al. 2017). Understanding both this correlation between fitness and dispersal as well as the distribution of effects for mutations impacting both of these characteristics is a major step forward in evolutionary biology, and could help us explain the different results already found in several experimental studies (Bosshard et al., 2017; Van Ditmarsch et al., 2013). Additionally, in natural systems this interaction between dispersal and load accumulation may explain important dynamics during colonization events. Given that invasive species present as ideal candidates for accumulation of expansion load in terms of rapid expansion and small founding population sizes, yet seem to exhibit no detrimental fitness effects, this mechanism of rescue and recovery due to increased dispersal could prove as an explanatory factor in their successful invasions as well as the successful spread of other natural range expansions (Arim et al., 2006; Hanski et al., 2002; Hughes et al., 2007; Lombaert et al., 2014; Monty and Mahy, 2010; Phillips et al., 2006; Simmons and Thomas, 2004; Szücs et al., 2017; Tayeh et al., 2013; Thomas et al., 2001). The improved understanding of the evolutionary forces and interactions between changes in fitness and dispersal ability will enhance our knowledge of what makes some species particularly successful at colonization, as well as what factors might contribute to formation of species range limits.

## Acknowledgments

We thank Ben Phillips for stimulating discussions on this subject, and Matteo Tomasini for helpful discussions about technical aspects of our study. KJG was supported by EMBO long-term fellowship ALTF2-2016.

## Online Appendix

### A Supplementary Figures

**Figure S1:**
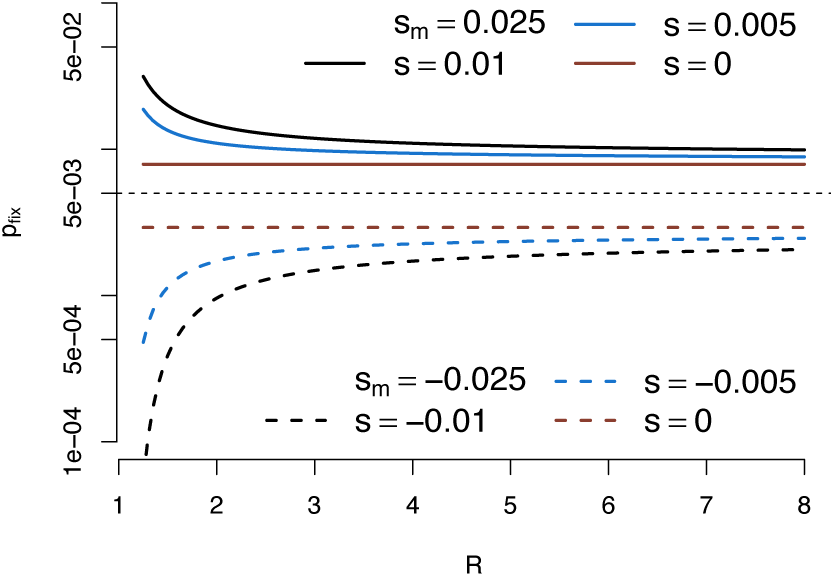
Fixation probability of pleiotropic mutations as a function of the population growth rate.

**Figure S2:**
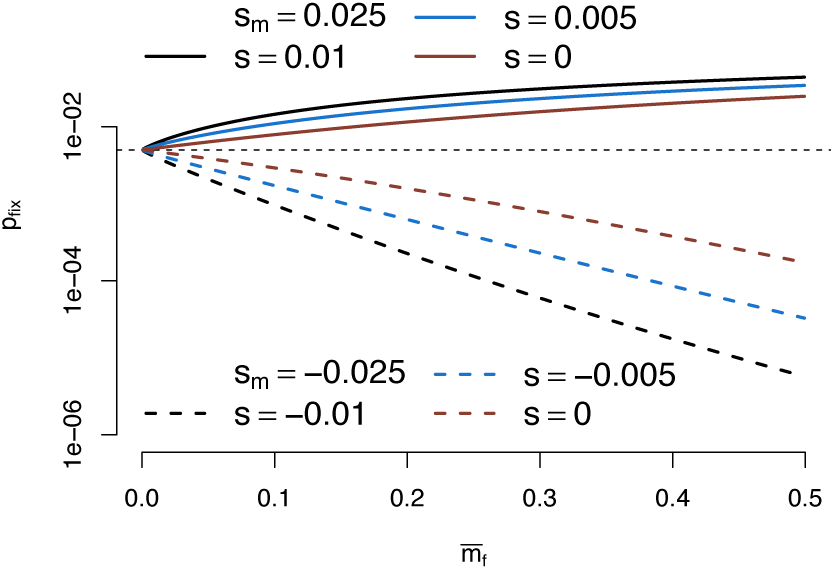
Fixation probability of pleiotropic mutations as a function of the mean migration rate at the expansion front.

**Figure S3:**
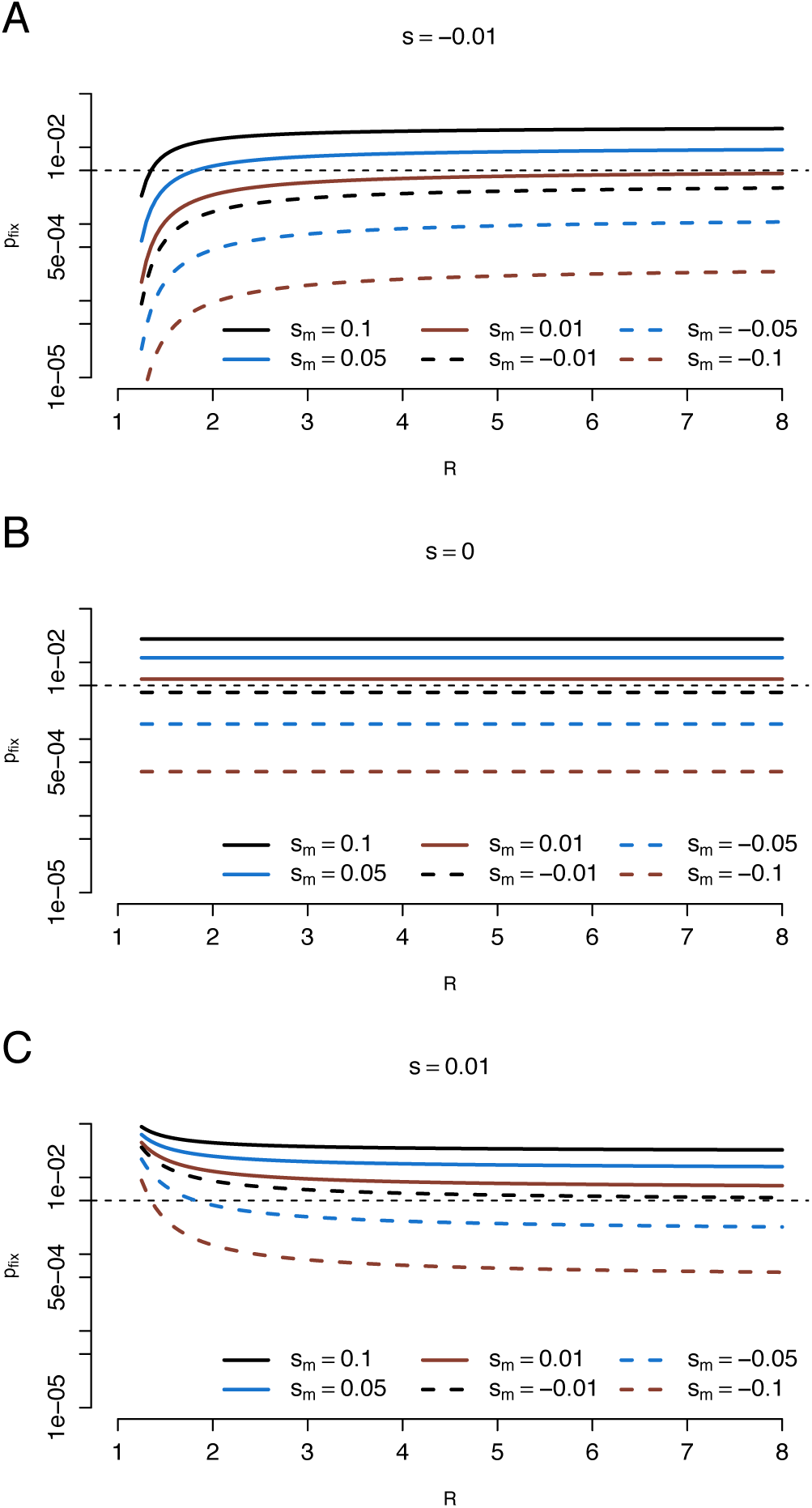
Fixation probability of pleiotropic mutations as a function of the population growth rate under varying scenarios of the mutational effect on fitness, *s*: deleterious (A), neutral (B), and beneficial (C).

### B Derivation of fixation probability

We use a diffusion approach to calculate the fixation probability of a mutation that affects fitness and/or migration rates. One “generation” in the diffusion approximation corresponds to the colonization of a single deme and starts just before a new propagule disperses. We consider a mutant that is present at frequency *p* when the population in deme *d*_*f*_ is at carrying capacity *K*. The expected frequency of the mutant in the propagule is then

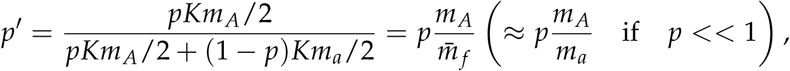

and the variance due to binomial sampling is

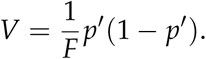

If the mutant’s frequency in the propagule is 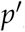, its expected frequency after the growth phase is

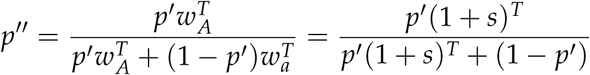

which is equal to

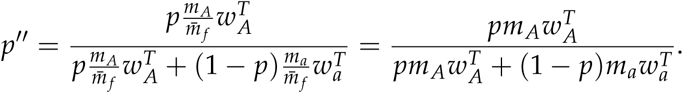

The expected change in allele frequency is then

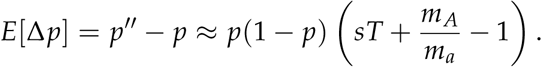

We assume that the stochastic sampling effects during colonization of a deme are the main contribution of genetic drift and therefore approximate the variance in allele frequency change by

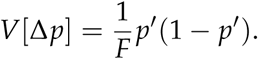

Next, we calculate the fixation probability using standard diffusion methods. The probability of fixation (conditioned on initial frequency *p*_0_) is given by:

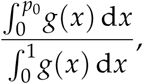

where

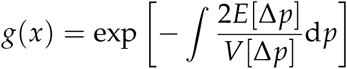

After some fundamental algebra we end up with

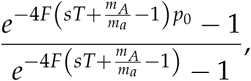

for the fixation probability, where *F* is the propagule size and *T* is the time until a deme is filled.

Note that this is Kimura’s (Kimura, 1962) original equation for the fixation probability

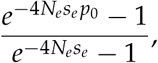

with effective selection coefficient

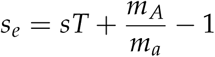

and effective population size *N*_*e*_ = *F*.

**Figure S4:**
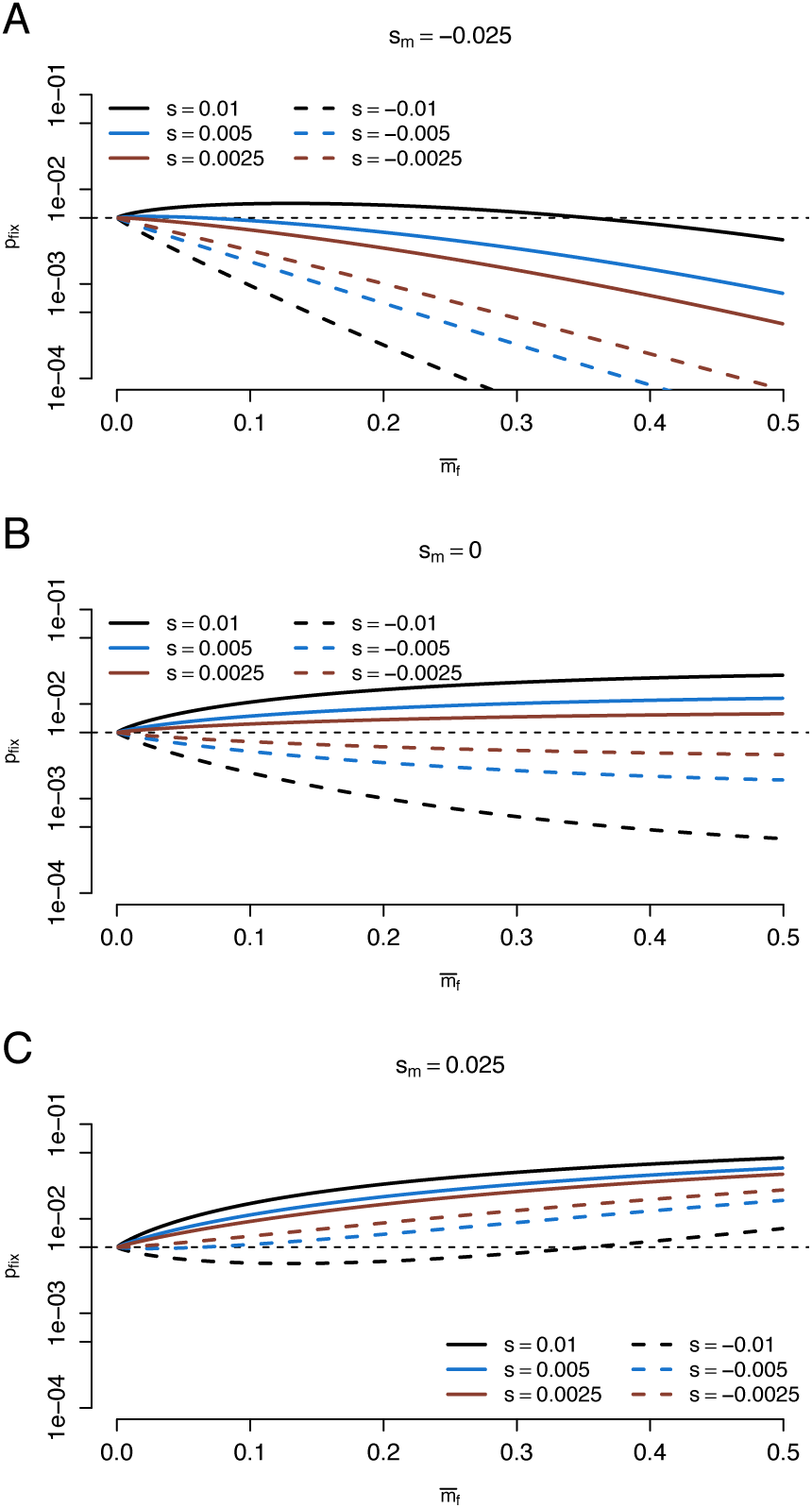
Fixation probability of pleiotropic mutations as a function of the mean migration rate at the expansion front.

### C Co-evolutionary dynamics

If mutations affect only fitness but not dispersal probabilities, Peischl et al. (2015) showed that the change in mean relative fitness at the expansion front can be approximated using the following equation

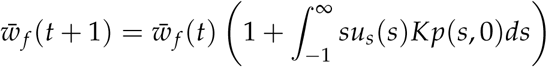

where *u*_*s*_(*s*) is the mutation rate of mutations with effect *s*. Here, we also assumed that mutations evolve independently of each other, that is, we ignore clonal interference. The parameter *F* measures genetic drift on the expansion front and *T* measures the length of the growth period during which selection occurs. Both *F* and *T* can depend on mean fitness, migration rates, and growth rates in models of hard selection (see Peischl et al. 2015 or Bosshard et al. 2017 for details). The integral in the equation calculates the expected long-term effect of each new incoming mutation, that is, whether they will be fixed or lost from the population at the expansion front, taking into account the effects of mutation rates, random genetic drift, natural selection, and spatial sorting. This model has been shown to be a good approximation for the evolution of mean fitness at the front of range expansions if the growth rate of populations at the front, *r*, is larger than the rate of gene flow, *m* (Peischl et al., 2013).

We next consider mutations that affect migration traits but have no effect on fitness. Let 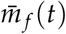 denote the mean migration rate of the population at the edge of the expansion at time *t*. The evolution of 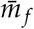 can be modelled analogously via

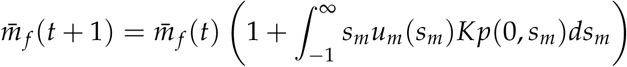

where *u*_*m*_(*s*_*m*_) is the mutation rate of migration modifier mutations that increase dispersal probabilities by a factor 1 + *s*_*m*_.

For pleiotropic mutations that affect both fitness and dispersal, the evolutionary dynamics of both traits follows

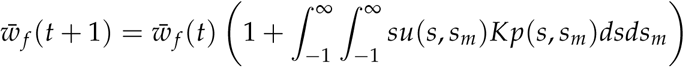

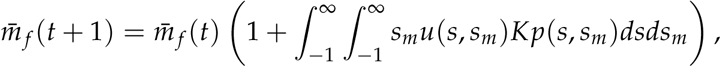

where *u*(*s, s*_*m*_) denotes the mutation rate of mutations with effect *s* on fitness and *s*_*m*_ on the dispersal probability. We assume that *s* and *s*_*m*_ are given by a bi-variate distribution with mean values 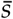 and 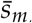, variances *V*_*s*_ and *V*_*m*_, and correlation *ρ* (e.g., a bi-variate Normal distribution). Switching to continuous time, the dynamics can be approximated by

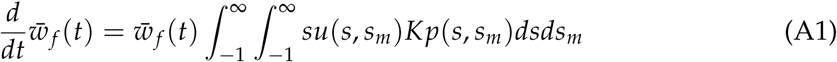

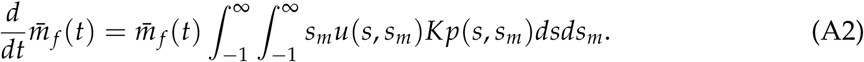

We define

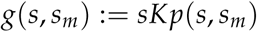

and approximate it by

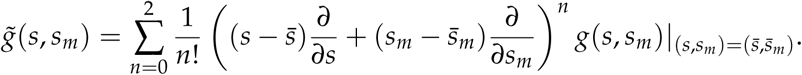

This allows us to approximate the integrals in equations (A1) by

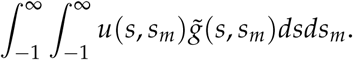

Equation (A2) can be treated analogously. These integrals can then be solved straightforwardly but the solutions are lengthy and uninformative and hence not shown here. Importantly, however, because 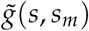 is quadratic in 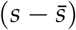 and 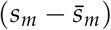, the contribution of the distribution of mutational effects *u*(*s, s*_*m*_) can be expressed solely in terms of the mean 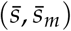, variances *V*_*s*_ and *V*_*m*_, and the correlation *ρ*. To gain a better intuition we proceed by further approximation. We re-scale the quantities *s*_*m*_, 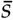, 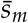, *V*_*s*_, *V*_*m*_ and *ρ* by *s*, and expand in a Taylor series around 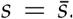. Ignoring thirdand higher-order terms in *s* (and switching back to the original variables) we can then approximate the dynamics of mean fitness and migration rate at the front by

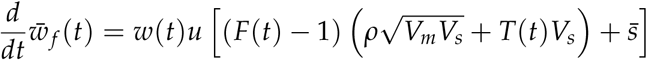

and

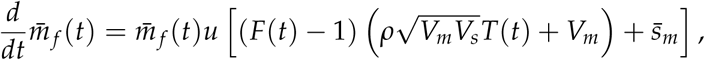

where 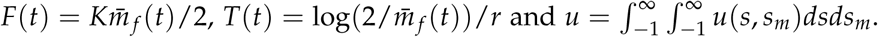.

If mutations affect either migration rates or fitness, but not both, we obtain

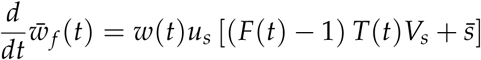

and

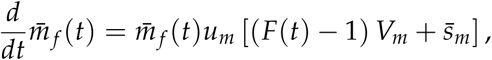

where *u*_*m*_ and *u*_*s*_ are the mutation rates of migration- and fitness-related mutations, respectively.

